# Autophagy induced accumulation of lipids in *pgrl1* and *pgr5* of *Chlamydomonas reinhardtii* under high light

**DOI:** 10.1101/2020.09.14.296244

**Authors:** Nisha Chouhan, Elsin Raju Devadasu, Ranay Mohan Yadav, Rajagopal Subramanyam

## Abstract

*Chlamydomonas (C) reinhardtii* cells (wild-type CC125 and *137AH*, and cyclic electron transport dependant mutants *pgrl1* and *pgr5*) were grown in high light 500 µmol photons m^−2^ s^−1^ where the growth was significantly enhanced after three days. The starch and lipid contents were also increased; however, starch content was decreased in *pgr5*. Further, the Nile Red fluorescence shows that a significant amount of lipid bodies were observed in *pgr5* cells under high light. Similarly, the electron micrographs show that large vacuoles were formed in high light stress despite the change in stacks of grana structure. We also observed increased production of reactive oxygen species (ROS) that could lead to autophagy. Inline, a significant increase of ATG8 protein was noticed in *pgr5*, which is a hallmark characteristic for autophagy formation. Consequently, the triacylglycerol (TAG) content was increased due to DGAT and PDAT enzymes’ expression, especially in *pgr5*. Here, the TAG synthesis would have been obtained from degraded membrane lipids in *pgr5*. Additionally, mono, polyunsaturated, and saturated fatty acids were identified more in the high light condition. Our study shows that the high light induces ROS, leads to autophagy and TAGs accumulation, which is stored as an energy source to acclimatize the algae.

## Introduction

Light is vital for expanding microalgae to perform efficient photosynthesis, carbon fixation, and rapid growth with high lipid productivity than plants. Microalgal species do not accumulate high amounts of neutral lipids under normal growth conditions. Fixation of CO_2_ by Calvin - Benson cycle, which primarily synthesizes carbohydrates, leads to the synthesis of lipids, which in turn, stored as lipid droplets, consists of high content of Triacylglycerols (TAG) under environmental stress conditions (Mondal *et al*., 2017). The biomass was increased while the acclimation of algae to high light stress (Yadav *et al*., 2020). Algal biomass is mostly composed of lipids, carbohydrates, and proteins. So, if there is an increase in lipid content, there will be a decrease in another component like proteins. Previous studies show it decreased photosynthetic activity and disorganization of thylakoid membranes in *C. reinhardtii* under high light (Nama *et al*., 2019). However, how light intensity affects the biochemical pathways were not studied adequately. It is imperative to see the changes in biochemical pathways under high light to observe biofuels’ most effective microalgae use.

An increase in light intensity produces reactive oxygen species (ROS), which regulate the autophagy mechanism. Autophagy is a process of stress-responsive mechanism, including nutrient stress, oxidative damage, and organelle deterioration (Liu and Bassham, 2012). This mechanism in Chlamydomonas regulates the degradative process in photosynthetic organisms. Also, it has divulged an essential role of autophagy in the control of lipid metabolism in algae. Inhibition of autophagy by the Target of Rapamycin (TOR) kinase showed by treatment of Chlamydomonas cells with the macrolide rapamycin resulted in increased ATG8 lipidation (Pérez-Pérez *et al*., 2010a) and vacuolization (Pérez-Pérez *et al*., 2010b). The previous results have been proven that ROS is an inducer of autophagy in algae (Pérez-Pérez *et al*., 2010a). High light stress induces photo-oxidative damage due to ROS production, which leads to the activation of autophagy in C. *reinhardtii* (Pérez-Pérez *et al*., 2012). A recent report has been shown that a starchless mutant of C. *reinhardtii* induces oxidative stress, which triggers autophagy that leads to TAG accumulation under nitrogen starvation (Pérez-Pérez *et al*., 2012). In a group of ATG (autophagy-related) proteins, the accumulation of ATG8 and ATG3 increased in a conditional repression line of a chloroplast protease ClpP1, suggesting the chloroplast proteolysis systems and autophagy are partially complement each other in Chlamydomonas (Ramundo *et al*., 2014).Additionally, these proteins were shown a similar function that of other organisms (Pérez-Pérez *et al*., 2010a; Pérez-Pérez *et al*., 2016). Loss of cytoplasmic structure with a significant increase in the volume occupied by lytic vacuoles and discharge of vacuolar hydrolases can be assigned as autophagic cell death markers (Van Doorn *et al*., 2011). However, the mechanism of autophagy is poorly understood in *C. reinhardtii* caused by high light stress.

Additionally, the neutral lipid accumulation can be achieved by exposing the cells to unfavourable conditions, such as removing the nutrients such as nitrogen, sulfur, iron, phosphate, salinity, and temperature (Alishah Aratboni *et al*., 2019). Most of the knowledge on TAG metabolism in *Chlamydomonas reinhardtii* has been gained from the N starvation (Siaut *et al*., 2011; Abida *et al*., 2015). Recent studies showed that cells exposed to small chemically active compounds could also accumulate TAG in *C. reinhardtii* (Wase *et al*., 2019). However, increase in light intensity influence the microalgal lipid production in Scenedesmus and accumulates more lipids with light intensities from 3000 to 6000 lux (Mandotra *et al*., 2016), also *Nanochloropisis* Sp at 700 µE m^−2^s^−1^ (Pal *et al*., 2011) and *Betyoccocusspe* at 6000 lux (Yeesang C *et al*., 2011). Further, *Haematococcus Pluvialis* (Zhekisheva *et al*., 2002), *Tichocarpus crinitus* (Khotimchenko and Yakovleva, 2005), and *Synechocystis* sp. (Cuellar-Bermudez *et al*., 2015) also showed an increased neutral lipid content under high light condition.

Lipid bodies are synthesized in algal chloroplast by fatty acid synthase complex. These newly synthesized free fatty acids are translocated to the endoplasmic reticulum (ER), where they convert Glyceraldehyde 3-phosphate (G3P) to Diacylglycerol (DAG) through various enzymes. DAG is converted to triacylglycerol by Diacylglycerol acyltransferases (DGAT) enzyme. So, the enzyme is considered to reinforce the assembly of lipids in microalgae. DGAT enzyme catalyzes the TAG synthesis pathway’s ultimate step, and phospholipid diacylglycerol acyl-transference (PGAT) also helps in accumulating TAG. However, it does not hook into the acyl CoA pathway. Here, PGAT transfer carboxylic acid moiety from a phospholipid to DAG to make TAG (Sorger and Daum, 2002; Shockey *et al*., 2006).

Currently, few studies have focused on increased fatty acid content with decreased photosynthetic activity, and retarded biomass ultimately inhibits microalgal growth in nutrient limitation. *Pgrl1* is required for efficient cyclic electron flow (CEF) and ATP supply for efficient photosynthesis (Tolleter et al., 2011; Leister and Shikanai, 2013; Shikanai *et al*., 2014; Steinbeck *et al*., 2015). A *pgr5* mutant has been characterized in Chlamydomonas (Johnson *et al*., 2014), which revealed that *PGR5* deficiency results in a diminished proton gradient across the thylakoid membrane accompanied by less effective CEF capacity. The CEF pathway has been suggested to provide ATP for lipid production during N starvation in *C. reinhardtii* mutants impaired within the proton gradient regulation 5 like 1 protein (*pgrl1*), which nonetheless accumulates significantly fewer neutral lipids under N starvation. However, under the high light condition, there is an increase in biomass and oil content after 3^rd^ day. So far, the accumulation of lipids has been well characterized in nutrient stress; however, this was not explored adequately under high light conditions. Since the biomass of C. *reinhardtii* is dependent on the solar light, which in turn converts into chemical energy using photosynthesis reactions, hence, *pgrl1* and *pgr5* mutants may play an important role, which might be due to an increase in stromal redox poise could lead to increased lipid production in *C. reinhardtii*.

In this study model, algae *C. reinhardtii* was propagated with different light conditions (50 and 500 µmol photons m^−2^s^−1^), to realize insights into the autophagy and lipid accumulation response to high light. Further, we examined the effect of light intensities on the biomass, cell ultrastructure with an abundance of starch and lipid droplets under high light-grown cells from wt, *prgl1*, and *pgr5. C. reinhardtii* cells also resulted in ROS accumulation, which in turn induce autophagy and lipid accumulation in *pgr5* when cells were grown in high light.

## Materials and methods

### Strains and culture conditions

The green microalgae *C. reinhardtii* wild-type (*wt*) strain *CC125* was obtained from the Chlamydomonas resource center (University of Minnesota) and *137AH*, mutants *pgrl1* and *pgr5* a kind of gift from Prof. Gilles Peltier, CEA-CNRS-Aix Marseille University, France and Prof. Michael Hippler, University of Munster, Germany. Cells were grown photoheterotrophically in Tris-acetate phosphate (TAP) medium at 25 °C with a photon flux density of 50 µmol photons m^−2^ s^−1^ in 250 mL conical flask shaken at 120 rpm in an orbital shaker at 25 °C. The seed cultures of *C. reinahrdtii* (*CC125, 137AH, prgl1*, and *prg5*) were harvested at an optical density (OD) of 0.8 and inoculated at OD of 0.2 for all the cultures and cultivated photoheterotrophically at 25 °C for 4th days at a continuous light intensity of 50 and 500 µmol photons m^−2^ s^−1^. Later the cultures were collected after 3^rd^ day with an OD of 0.8 at 750nm. All the experiments were conducted with this condition. Light intensity was measured and adjusted with a light meter (Hansatech).

### Biomass determination

In order to calculate the total biomass of cells (*CC125* and *137AH*, and mutant *pgrl1* and *pgr5*), 5 mL of culture was collected after 3^rd^ day of growth in high light along with control by centrifugation at 1000 x *g* for 10 min at RT, and the supernatant was discarded. The cell pellet was washed with distilled water transferred into a pre-weighed 15 mL falcon tube. The excess media was removed by short centrifugation. Tubes containing cell pellet was kept at −109 °C for 12 h. Total dry weight was quantified by subtracting the empty tube.

### ROS detection by confocal microscopy

The total ROS was detected with 2,7dichlorodihydrofluorescein diacetate (H_2_DCFDA), a fluorescence dye to detect the ROS in live Chlamydomonas cells. Cells at a 3 million density were collected from all the conditions, and H_2_DCFDA staining was performed at 10 µM concentration (Upadhyaya *et al*., 2018). Further, cells were incubated with dye for 1h at RT in a continuously rotating shaker under dark. Images were captured using Carl Zeiss NL0 710 Confocal microscope. H2DCFDA was detected in 500 to 530 nm bandpass optical filter with an excitation wavelength of 492 nm and an emission wavelength of 525 nm. Chlorophyll auto-fluorescence was detected using an optical filter of 600 nm. Samples were viewed with a 60x oil immersion lens objective by using the ZEN 2010 software.

### Localization of ATG8 from immunofluorescence microscopy

For immunofluorescence, cells were fixed in the solution consisting of 4% paraformaldehyde and 15% sucrose dissolved in phosphate saline buffer (PBS) for 1h at RT and later, the cells were washed twice with PBS buffer. Cells were permeabilized by incubation in 0.01% Triton X100 in PBS for 5 min at RT and washed twice in PBS. The samples were transferred to sterile Eppendorf tubes and blocked with a 1% BSA (w/v) in PBS for 1 h. Samples were incubated with anti-ATG8 diluted (1:250) in PBS buffer, pH= 7.2 containing 1% BSA overnight at 4 °C on a rotatory shaker. Cells were then washed twice with PBS for 10 min at 25 °C, followed by incubation in a 1:200 dilution of the fluorescein Dylight 405 labelled goat anti-rabbit secondary antibody (Sigma) in PBS-BSA for 2 h at 25 °C. Finally, cells were washed three times with PBS for 5 min. Images were captured with Carl Zeiss NL0 710 Confocal microscope. ATG8 localization was detected with excitation of Dylight 405 nm and emission at 420 nm by analyzing the images with ZEN software.

### Lipid body analysis by Nile Red staining

The impact of high light on non-polar lipid accumulation was studied with confocal microscopy. Cells were collected after 3^rd^ day growth in high light along with control. To stop the cells’ mobility, 5 µL Iodine solution (0.25 g in ethanol (95%)) was mixed and kept for 5 min under dark at RT. The cell suspension was then stained with NR (1 µg mL^−1^ final concentration, Sigma) followed by 20min incubation in the dark. After staining, the samples were placed in a glass slide with a coverslip. Images were captured using Carl Zeiss NL0 710 confocal microscope. For the detection of neutral lipids, a 488 nm scanning laser was used with a 560 to 615 nm filter. Samples were viewed with a 60x oil immersion lens objective, and the ZEN 2010 software package was used for image analysis.

### Nile Red visible light assay for triacylglycerol

The *C. reinhardtii* cultures were diluted to a density of 3.0 × 10^6^ cells mL^−1^ and placed within 96 well microplates wells. To every well, a 5 µL aliquot of 50 µg mL^−1^ NR; (Sigma; ready in acetone) was added, and after thorough mix, the plate was incubated in the dark for 20 min. The neutral lipids were then measured by visible light at a 485-nm excitation filter and a 595-nm emission filter employing a plate reader (Tecan infinite M200, Magellan). Quantification was achieved by employing a standard curve ready with the lipid Triolein (Sigma T4792).

### Flow cytometry measurements

Flow cytometer Cali bur (BD Falcon, USA) was used for lipid analysis by cell sorting. Cells were stain with 1 µg/mL NR (100 µg/mL stock in acetone) in the dark for 30 min before flow cytometry analysis. Measurement of 10,000 cells per sample was analyzed and used for sorting. The fluorescence reading was obtained at a 488-nm excitation filter and a 545-nm emission. The distribution of the cells stained with NR was expressed as a percent of the total number of cells using the software Flow Jo and mean fluorescence was calculated.

### Total chlorophyll and starch quantification

*C. reinhardtii* cells were harvested by centrifugation (1000x *g* for 10 min) after 3^rd^ day at a cell density of 3.0 × 10^6^. Cell pellets were resuspended in methanol for chlorophyll extractions, and the supernatant was used to measure the total chlorophyll, according to Porra et al. (Porra *et al*., 1989). Total starch quantification was quantified by an Anthrone method (plate reader assay). For quantification of starch, the cell pellet was resuspended in 50 µL Milli-Q water and added 150µL of Anthrone chemical agent (0.1g in 100mL of conc. H_2_SO_4_ 98%) to the 50µL algal cells. Initially, plates were incubated at 4 °C for 10 min and then incubated at 100 °C for 20 min. Later on, plates were placed at temperature until color development. Plates were measured at 620 nm on an assay plate reader (Teccan M250), and total starch was calculated according to Wase et al. (2019).

### Lipid extraction and thin layer chromatography analysis

For lipid extraction, 5-mg of lyophilized dry biomass was weighed equally from all the conditions. Total lipids were extracted as described by Bligh and Dyer (Bligh and Dyer, 1959). Cell powder was resuspended in a mixture of methyl alcohol and chloroform (2:1) and 0.9% KCl. Vortex for 5 min and centrifuged at 3000× *g* for 5 min for phase separation. The organic layer (lower layer phase) was transferred to a clean Eppendorf tube. The solvent was evaporated by purging nitrogen gas. The dried extract was then dissolved in 50 µL chloroform. The polar lipids were analysed by two-dimensional thin layer chromatography (TLC), the first dimension lipids were separated with a solvent mixture consisting (chloroform: methanol: water), and after air dry, the second dimension was done with a solvent mixture containing (chloroform: methanol: glacial acetic acid: H_2_O). Later on, all the lipids were stained with Phosphomolybdic acid (Sigma) dissolved in 95% crude ethanol.

### Immunoblots

Cells were collected by centrifugation. The cell pellet was solubilized in protein extraction buffer (0.1M DDT, 4% SDS, and 0.1 M Tris (pH-7.6), and all the samples were kept for heating at 95 °C for 5 min. Cell suspensions were centrifuged at 2810x *g* for 10 min, and the supernatant was collected. Protein concentration was quantified by using the Bradford method and 5 µg of proteins was loaded per well. The individual proteins were separated with 15% Bis-Tris gels for DGAT2 and PDAT1 proteins and 12% for ATG8 and Histone3 (H3) proteins. Separated proteins were then transferred to a nitrocellulose membrane, according to Twobin *et al*. (1979). Specific primary antibodies were purchased from Agrisera (Sweden) and used at dilutions of DGAT2A (1:2500) and PDAT (1:2500). H3 (1:10,000) was used to see equal loading. The secondary antibody anti-IgG raised from Goat (1:10,000) was used for detection. Immunoblots were obtained by the Chemi-Doc touch imaging system (Bio-Rad) by using Chemiluminescence.

### FAME analysis

The 5 mg lyophilized sample was dissolved in 1000 µL of acetonitrile, 0.1% formic acid (v/v), and vortexed for 1h at 900 rpm at room temperature. The solution was then centrifuged at 13000 rpm 10 min at 4 °C. 100 µL of supernatant was collected and dried the pellet using a speed vacuum. The pellet is dissolved in 90 µL of acetonitrile 0.1% formic acid (v/v) and spiked with 10 µL of the standard internal mixture (Table. 1). The solution is vortexed for 30 min at 750 rpm at RT and speed vacuumed to dry at 40 °C. The extracted lipids were derivatized with n-Butanol for 20 min at 60 °C, then again speed vacuumed to dry at 60 °C for 15 min. The dried pellets were dissolved in 100 µL of ACN 0.1% FA and vortexed for 5 min at 750 rpm on a thermomixer. The samples were centrifuged at 13000 rpm for 10 min at RT. The supernatant was taken into HPLC sample vails for further analysis. The 100 µL extracted lipid samples were brought to RT and filled into individual HPLC vails with 200 µL inserts and placed into the autosampler of Nexera X2 UFLC, connected to LC-MS/MS (SHIMADZU 8045) MS system with an ESI source. The data was analyzed by Lab solutions software-the concentrations calculated by measuring the area under the curve (AUC) for the internal standards.

## Results

### Growth and pigment analysis under high light

The *CC125* and *137AH, pgrl1*, and *pgr5* (cyclic electron transport dependant mutants) were grown in TAP medium at light intensities of 50 (as a control) and 500 µmol photon m^−2^s^−1^ for 4th days. Under 50 µmol photon m^−2^s^−1^ light intensity, all the cultures were grown usually. At 500 µmol photon m^−2^s^−1^ light condition, their growth was reduced to half, especially in mutants (Fig. 1). The growth of *CC125* and *137AH* cultures increased, but *pgrl1* and *prg5* mutant exhibited slower growth in high light conditions after two days (Fig. 1A). Our results showed that the cell growth pattern of the *CC125* and *137AH* were increased 25 and 30% whereas *prgl1* and *pgr5* growth were decreased 21 and 30% compared to the WT under high light (500 µmol photon m^−2^s^−1^). However, these mutants growth is still higher than the 50 µmol photon m^−2^s^−1^ light. Mainly, *pgr5* mutant shows impaired growth, whereas *pgrl1* was not affected under the same high light condition (500 µmol photon m^−2^s^−1^) after 3^rd^ day of growth (Fig. 1A). *CC125, 137AH*, and *pgrl1* showed exponential growth, while *pgr5* shows slower growth under high light intensity after 3rd days. We have studied the effect of light intensity on biomass; the *CC125, 137AH*, and *pgrl1* show increased (twofold) in yield of biomass under average and high light intensities. In contrast, *pgr5* showed decreased (half of its normal condition) biomass after 3^rd^ day (Fig. 1C).

**Fig. 1.**
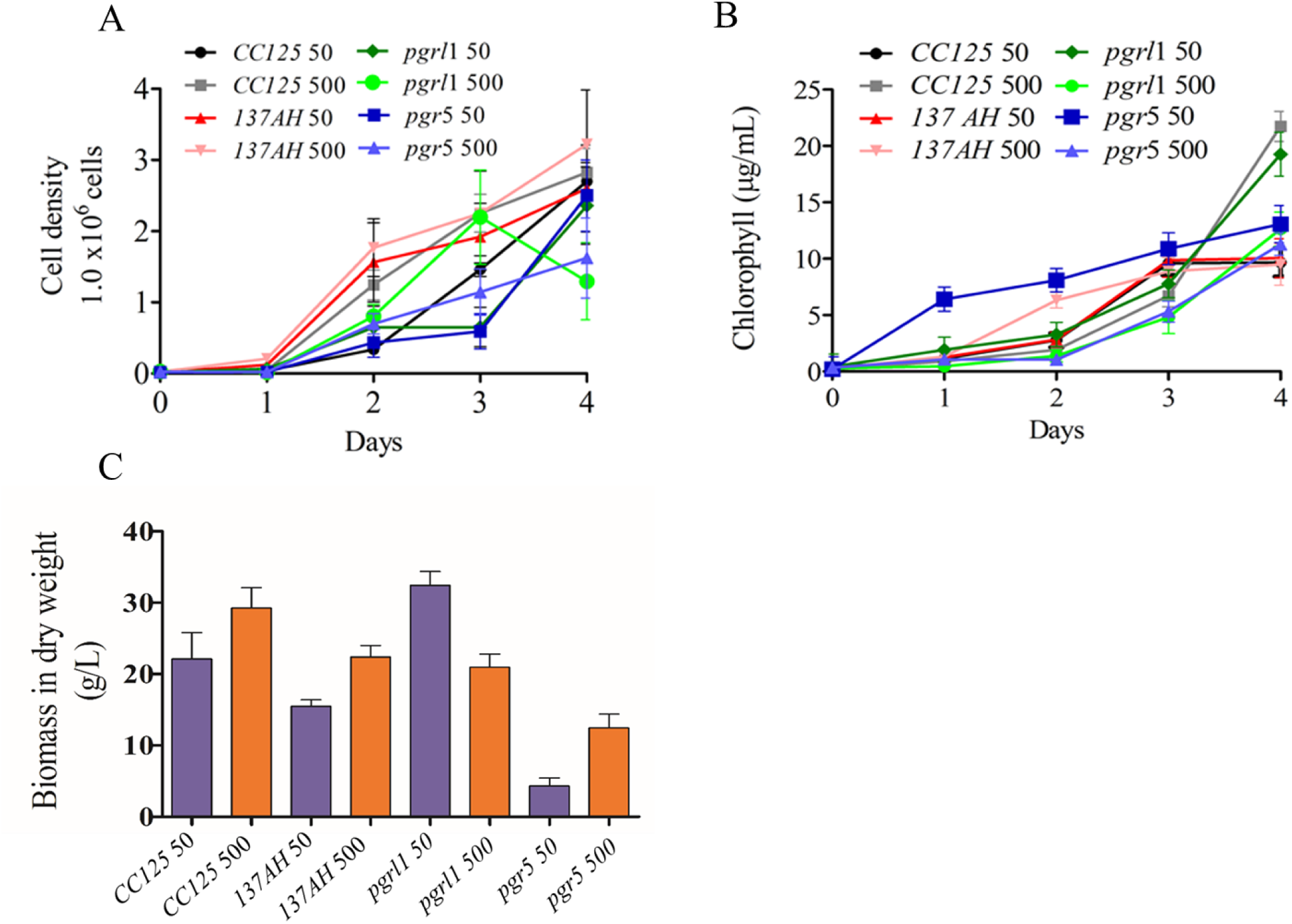
Cell growth, total chlorophyll, and dry biomass were monitored from cells grown under normal light as well as high light conditions (A) cell growth was analyzed by cell counting by hemocytometer collected for every 24h of *CC125, 137AH, pgrl1*, and *pgr5* from 50 µmol^−2^ s^−1^ and 500 µmol photons m^−2^s^−1^ conditions. (P<0.050). (B) Chlorophyll content was quantified from the conditions mentioned above with different time intervals of 24h, 48h, and 72h (P < 0.004). (C) Total biomass in dry weight was calculated after 3^rd^ day of growth of cells (*CC125* and *137AH, pgrl1* and, *pgr5*) grown from light conditions (P < 0.004). All the experiments were done with three independent times, and error bars represent the mean ± SD (n=3).

To determine the effect of high light on the pigment level associated with Chlamydomonas’ photosynthetic complex, we have measured the chlorophyll and carotenoid content from standard and high light-grown cells. Chlorophyll content was significantly reduced to 50% in *pgrl1* and *pgr5* mutant, respectively, under high light intensity. At the same time, we observed that increased carotenoid content in *pgrl1* (15%) and *pgr5* (20%) (Fig. S1). It is known that increased carotenoid content acts as a protective mechanism against light stress in the Chlamydomonas (Li. Z *et al*., 2009). Carotenoids under high light conditions generally tend to increase in order to protect PSII from photoinhibition. Data shows that an increase in light intensity results in increased specific growth rate, increased carotenoid content, and biomass production of *CC125, 137AH*, and *pgrl1* strains except *for pgr5*. We assume that decreased growth might be due to ROS production, as shown in Fig. 2. Our results have elicited us to check further studies and detect the ROS in high light stress.

**Fig. 2.**
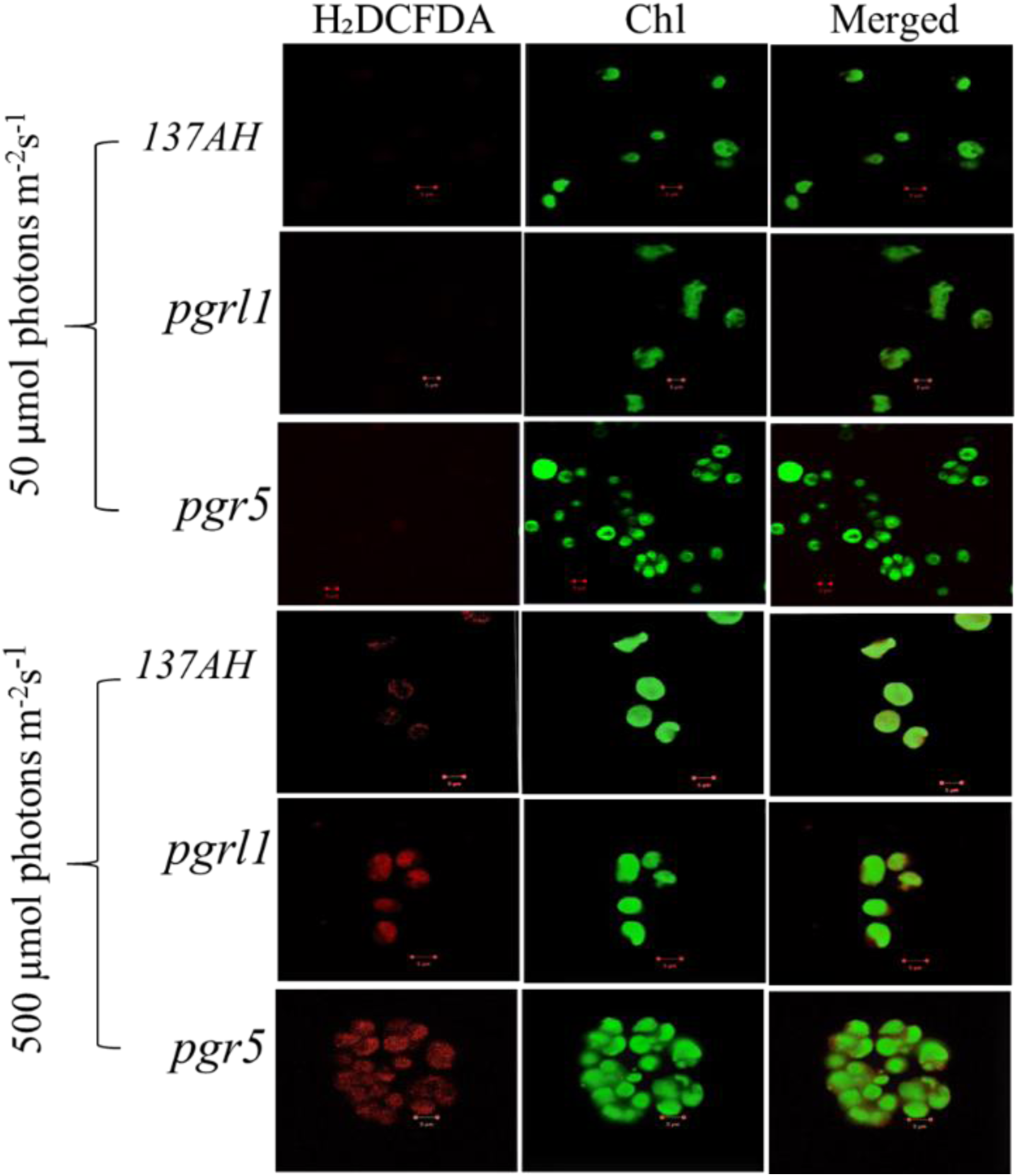
Chlamydomonas cells were collected from the mid-log phase grown under normal and high light for 3^rd^ day. They measured ROS using H2DCFDA (10 µM) staining, and cells were imaged using the Zeiss 510 confocal microscope. The images were collected with three individual measurements for all the conditions. Three independent experiments were conducted (n=3) and analyzed with Zeiss software.

### ROS induction in high light

In order to determine oxidative stress, cells were stained with H_2_DCFDA. ROS was observed under confocal microscopy, where an increase in ROS was observed under the high light condition in *CC125, 137AH, prgl1*, and *pgr5*. However, this increase was predominant in *pgr5* (Fig. 2). The generation of more ROS in *pgr5* could probably explain the reduced growth rate. Further, these results suggest that ROS formation under high light may contribute to autophagy and lipid accumulation. Reports show that ROS may induce multiple metabolic functions like lipid metabolism and autophagy (Shi *et al*., 2017). High light is one of the factors responsible for ROS formation because of over reduction of photosynthetic electron transport chain, which might generate ROS that induces autophagy in Chlamydomonas (Pérez-Pérez *et al*., 2012). Based on the above reports, we have conducted experiments related to autophagy induced by high light.

### Protein analysis by immunoblot

We have focused on the enzymes which are involved in the TAG synthesis known as the Kenndey pathway. This pathway is mainly catalyzed by Acylation of DAGs by DGAT and PDAT enzyme in which acyl-CoA as a substrate to form TAG in the endoplasmic reticulum. To determine the involvement of DGAT and PDAT in high light response in *C. reinhardtii* and their protein level regulation was determined (Fig. 3). The PDAT and DGAT were transiently upregulated in response to high light conditions. Protein expression achieved the maximum level after 3^rd^ day, specifically in *pgr5* expression of PDAT compared to *pgrl1* (Fig. 3). Our results confirm an increase in TAG in *pgrl1* and *pgr5* because of the increase of two enzymes. As there is more accumulation of TAG in the *pgr5* mutant, it could reduce membrane lipids. That was in agreement with the electron microscopic data that the stacks of thylakoids disturbed, which could be changed in membrane lipids (Fig. 6). As it is already proven that a decrease in MGDG is usually lead to TAG acceleration under stress condition. In vascular plants, a drastic decrease in MGDG content occurs during freezing or ozone treatment (Li *et al*., 2008). Our group recently reported that TAG accumulation originated from the degradation of MGDG under Fe deficiency from C. *reinhardtii* (Devadasu *et al*., 2019).

**Fig. 3.**
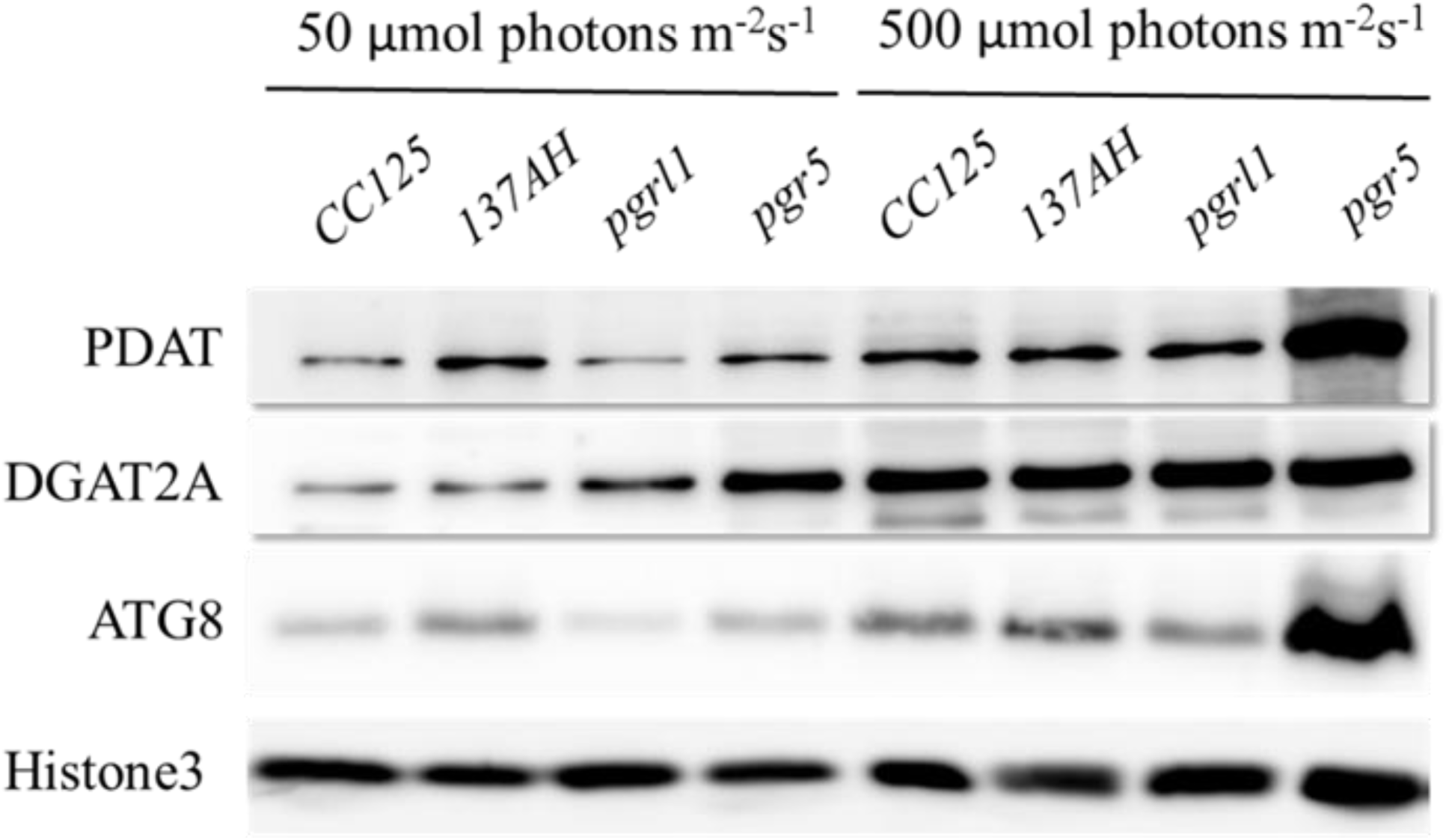
Protein identification from western blot analysis. The proteins were separated by denaturing gel electrophoresis, transferred to nitrocellulose membranes, and probed for the indicated proteins. Western blot analysis of DGAT2A (Acyl-CoA: Diacylglycerol acyltransferase) and PDAT1 (phospholipid diacylglycerol acyl-transference) evaluated from 3^rd^-day cells under the average and high light condition and each lane (5µg) protein was loaded on to 10% Bis-Tris gel in denaturing condition; ATG8 protein was resolved on 15% Bis-Tris gel in denaturing condition. Histone (H3) was used as a loading control. All the blots were in three independent times (n=3) and obtained similar results.

In order to assess all the membrane lipids of C. *reinhardtii*, 2-D thin chromatography was carried out. Thin-layer chromatography plates were stained by phosphomolybdic acid hydrate and further observed reduced in membrane lipids under high light conditions, especially in *pgr5* mutant (Fig. S2). Here, we detected the accumulation of TAG in *C. reinhardtii* under high light, is accompanied by substantial degradation of chloroplast membranes. Thus fatty acids from the chloroplasts and other intracellular membrane systems might have converted into TAG.

### Cellular localization of ATG8

We have used an anti -ATG8 antibody to examine this protein’s cellular localization in immunofluorescence microscopy (Fig. 4). ATG8 proteins localize as small red dots in the cytoplasm under the normal light condition in *137AH, pgrl1*, and *pgr5* mutant. However, under high light conditions, number and spot size significantly increased (50%) in *pgrl1* and (70%) in *pgr5* mutants, which shows activation of autophagy and formation of autophagosomes close agreement with high accumulation of ATG8 detected by western blotting. However, the ATG8 expression is predominant in *pgr5* under high light conditions. Previous reports have shown that oxidative stress-induced the autophagy in Chlamydomonas cells (Pérez-Pérez *et al*., 2012; Perez-Martin *et al*., 2014). These results suggested to us more ROS due to high light, and it may induce autophagy in *pgr5* than *137AH* and *pgrl1*. It induces autophagy due to oxidative stress, and it may trigger the lipid metabolism in Chlamydomonas cells. Further, we have carried out experiments related to lipid analysis.

**Fig. 4.**
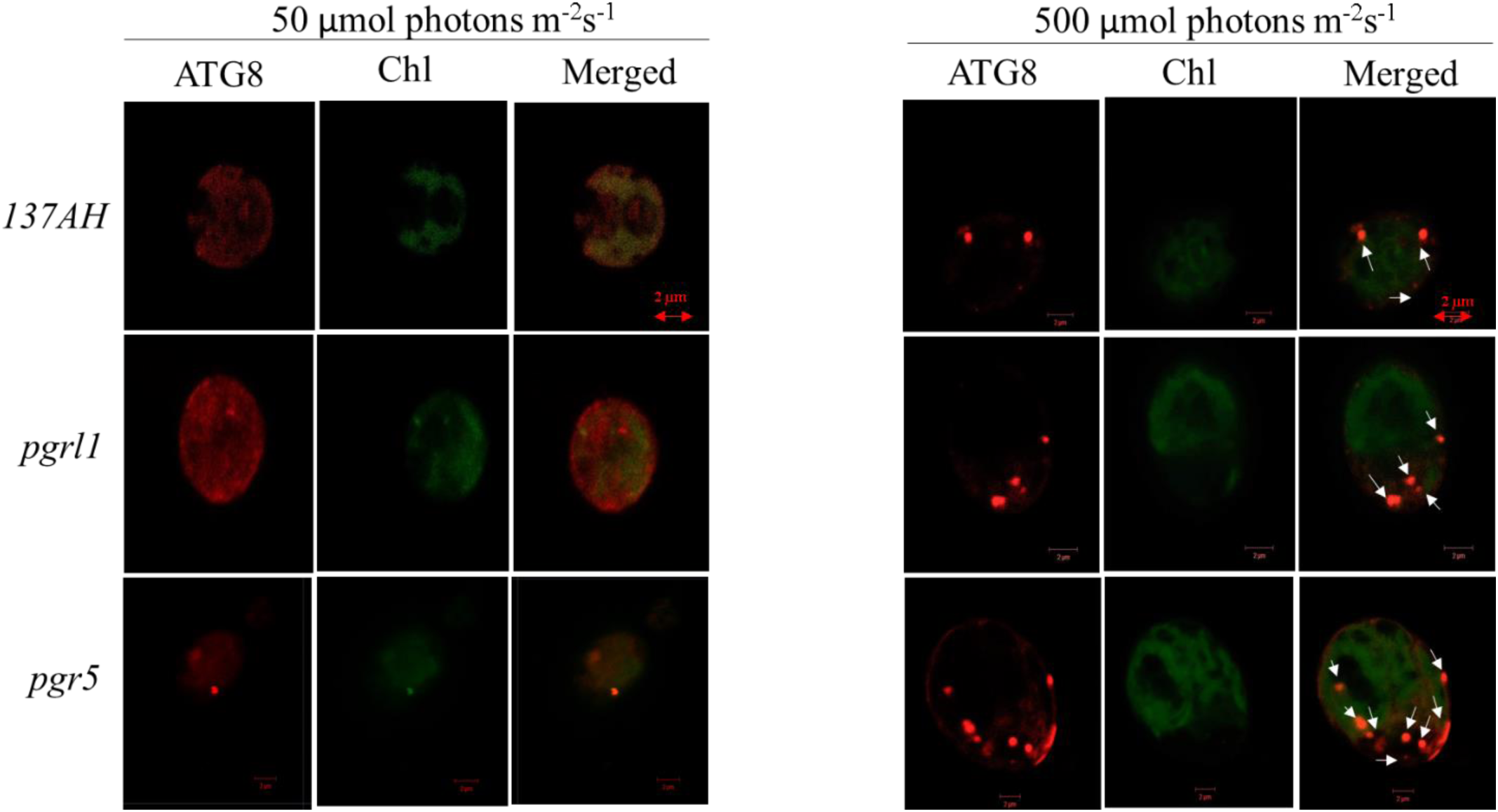
ATG8 accumulation in *C. reinhardtii* cells under high light condition. Autophagy induction in *C. reinhardtii* cells collected at phase cells (3.0 × 10^6^ cells) grown in TAP medium under continuous light of 50 and 500 µmol photons m^−2^s^−1^. Samples were collected after 3^rd^ day and immunoassay with anti-ATG8 antibody (in red). Chloroplasts were visualized by auto-fluorescence (in green). The image was taken using a Zeiss confocal microscope.

### Lipid droplet analysis from confocal microscopy

We have stained the cells with NR for lipid studies, which specifically binds to lipid bodies in the cell and gives a characteristic yellow-orange fluorescence. Our results clearly show the overall increase (40%) of lipid droplets under high light in *wt, pgrl1*, and *pgr5* mutants (Fig. 5A). However, autofluorescence due to chlorophyll was less in *pgrl1* and *pgr5* than control, which indicates that decreased chlorophyll content under high light, even though cells had a significant accumulation of lipid bodies (Fig. 5A). Although lipid body formation’s greatest density appears in *pgrl1* and *pgr5*, lipid accumulation was observed in all strains under high light conditions. However, lipids accumulation is significant (60%) in *pgr5* compared to other strains under high light. We have also tested NR fluorescence from FACS to assess the lipid accumulation in the high light conditions. We obtained two-fold mean fluorescence intensity from all the strains with high light conditions than normal light conditions (Fig. 5B). Our results suggested that high light induce the lipid metabolic pathways, thereby it led to the accumulation of triacylglycerol in the cell.

**Fig. 5.**
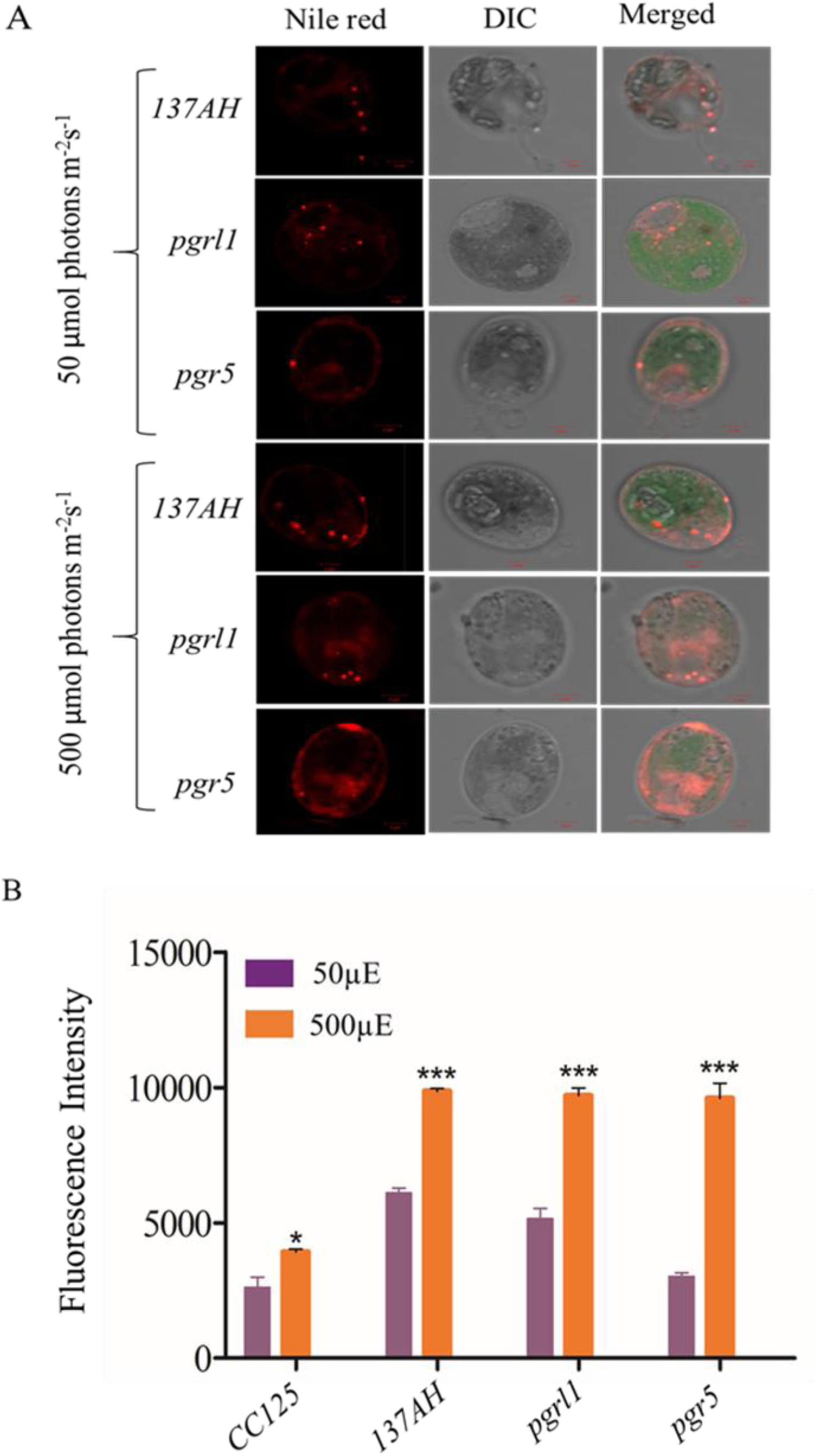
Lipid droplets were identified through confocal microscopy and FACS analysis. (A) Cells grown under light (50 µmol photons m^−2^s^−1^ & 500 µmol photons m^−2^s^−1^) conditions were stained with Nile Red (5 µM/mL) for lipid droplets in *C.reihnardtti* strains, *CC125, 137AH, prgl1* and *pgr5*. The images were collected with three individual measurements for all the conditions. (B) FACS analysis was carried out for lipid accumulation with BD Fertessa, USA. All the measurements were done with the PE-A filter. The analysis showed a histogram and mean fluorescence value for comparison from normal light to high light condition. An unstained condition was kept to minimize the chlorophyll autofluorescence from cells. Cells were stained with Nile Red, and measurements were done after incubated for 20 min under dark at RT. Statistical analysis and mean fluorescence values were represented with each condition. The measurements were repeated with three individual cultures, and error bars represented the mean ± SD (n=3). Samples were done with student *t*-test, and it significantly accepted (P < 0.004).

**Fig. 6.**
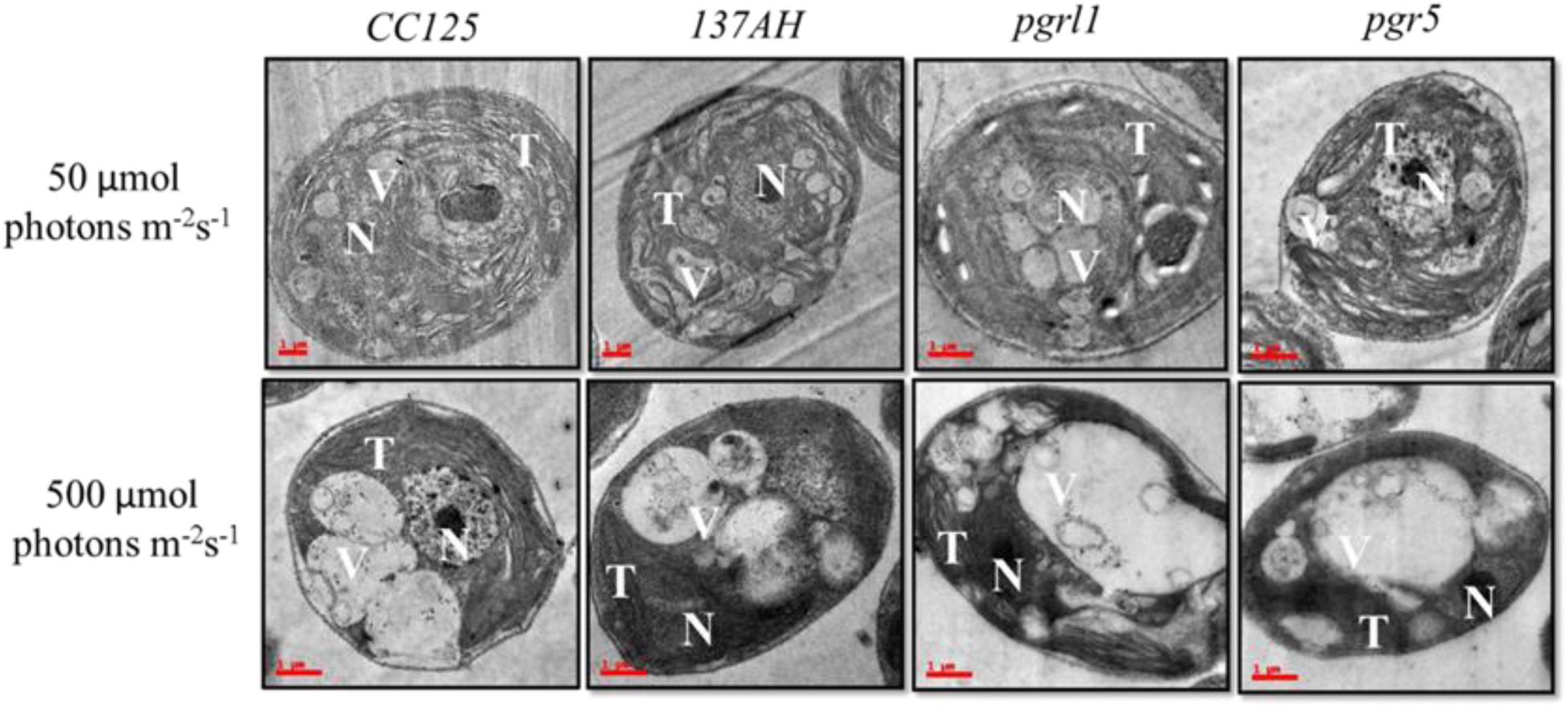
Ultrastructure analysis of Chlamydomonas cells during normal (50 µmol photons m^−2^s^−1^) and high light (500 µmol photons m^−2^s^−1^). Transmission electron micrographs of *C. reinhardtii* cells control *CC125* and *137AH* and mutants *pgrl1* and *pgr5* grown under photoheterotrophic conditions under normal and high light. Representative images for the cells sampled after 3^rd^ day, (A and B) N-nucleus; O, T-thylakoid membranes; V-vacuole. Three independent experiments were conducted from each sample (n=3).

### Neutral lipid identification by NR fluorescence

The neutral lipids were semi-quantified by using NR fluorescence from an equal number of cells. NR fluorescence results show that the neutral lipid level was increased with equal cell density. NR fluorescence of Chlamydomonas cells *CC125, 137AH, pgrl1*, and *pgr5*, were calculated at every 24 h while growing them in light at 50 and 500 µmol photon m^−2^s^−1^, as shown in Fig. S3. After 24 h, cells did not cause any change in total lipid content. However, after 48 h of high light growth, there was an increase in NR fluorescence. In contrast, there is an increase in signal at 750 nm, suggesting that cells are still healthy with increased biomass. Overall, high light-grown cells exhibit a relative two-fold increase in neutral lipid content over 3 to 4 days. However, this accumulation of neutral lipids is predominantly more in *pgr5* mutant when compared to other strains. Therefore, our results indicates that an increase in light intensity not only increases the biomass but also facilitates the production of neutral lipids in Chlamydomonas. Further, we have performed electron microscopic studies to determine whether high light induces the lipid accumulation in the cells.

### Transmission electron microscopy studies

To see the clear lipid vacuoles in the cell, we have carried out the transmission electron microscopy (TEM). It reveals that *prgl1* and *pgr5* exhibits a large cytoplasmic vesicle under high light grown cells (Fig. 6). It also revealed that the lipid composition’s membrane structure is aberrant under high light, especially the *pgrl1* and *pgr5* mutants. The majority of the thylakoid membranes were disrupted in high light grown cells. While in the wild type cells in control, the thylakoid membranes are arranged as layer (stacked) in the chloroplast, with some membrane appressed. Recently we reported that photosynthetic efficiency reduced under the high light condition, which may be due to change in chloroplast structure (Yadav *et al*., 2020). Such phenotype raised the question that mutant strains are defective in generating the functional membrane under high light conditions. It has been already reported that autophagic bodies accumulate in the vacuoles of plant cells under Concanamycin A treatment because vacuolar hydrolase cannot act (Thompson *et al*., 2005). However, in our case, under the high light condition without any inhibitors, we have observed a high degree of vacuolization and an increase of vacuole size in mutants. It has also been reported that vacuole lytic function is needed to synthesize TAG and lipid bodies in Chlamydomonas cells when subjected to light stress (Couso *et al*., 2018). Further, we have observed the large starch granules in the cell, we have tested the total starch from all the conditions.

### Starch assay

The starch content was quantified in the normal light condition of *CC125, 137AH* and *pgrl1* strains showed similar starch contents as 18 µg, 10 µg, and 17 µg starch/10^6^ cells, respectively. While under the highlight condition, the starch content of *CC125, 137AH*, and *pgrl1* was doubled, which shows a significant increase in starch content. The starch content was cross-checked with a starch deficient mutant (*sta6*) with normal light and under high light. It shows that 8 µg of total starch/10^6^ cells in both standard and high light conditions (Fig. 7). Overall, our reports indicate that starch content has increased under the high light condition in wild type and *pgrl1*. However, it reduced in *pgr5* mutant under high conditions. Environmental stress, like temperature and light, induces the starch, leading to the synthesis of lipids in green algae. In order to survive under unfavourable conditions, microalgae change their metabolism and convert excess energy into an energy-rich product such as lipid or starch, or proteins (Heredia-Arroyo *et al*., 2010). However, *pgr5* may have an altered NADPH and ATP ratio during photosynthesis, which may switch metabolic pathways that lead to lipid accumulation. These results indicate that high light is inducing the lipid pathways due to more energy equivalents in the *pgr*5 mutant. Further, we have tested the protein/enzyme of lipid pathway from all the conditions.

**Fig. 7.**
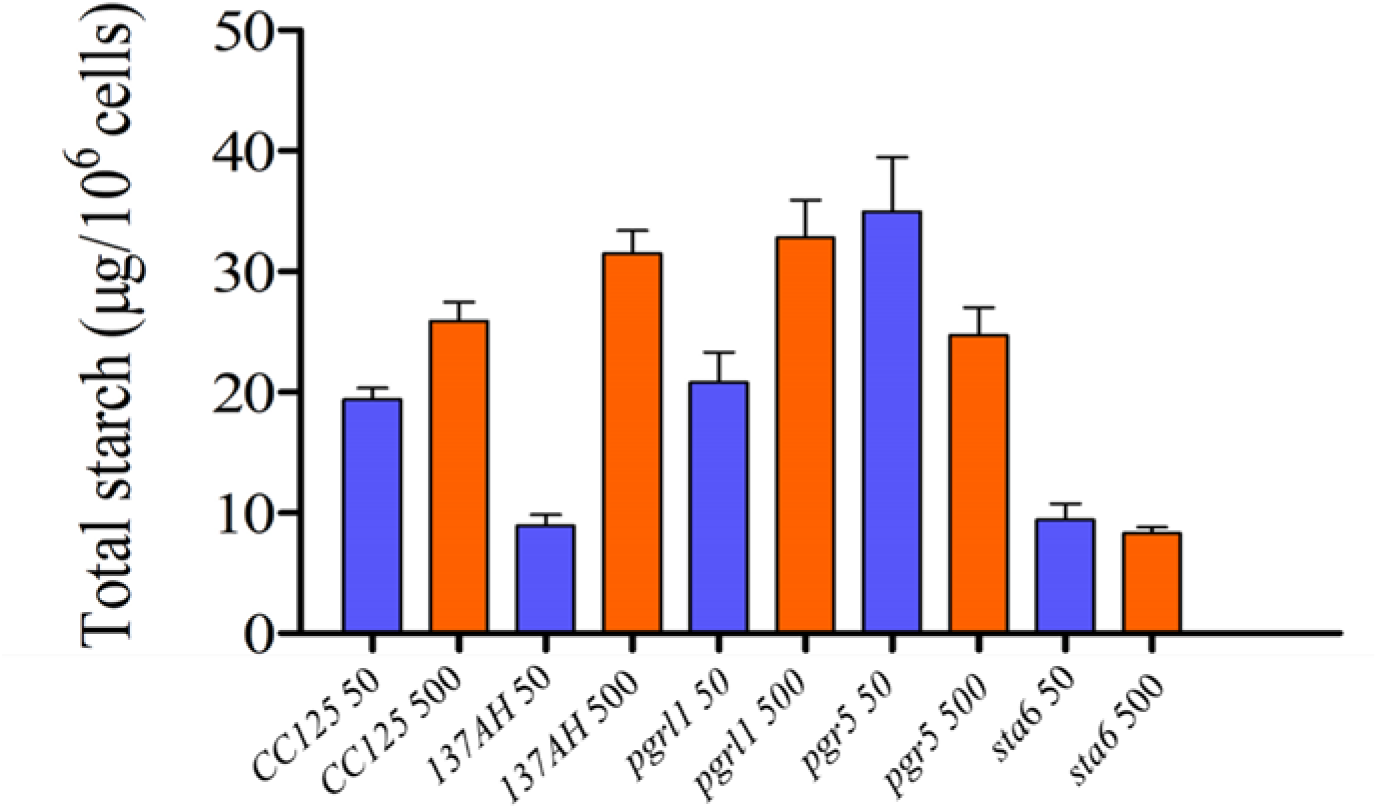
Total starch content quantified from the cells grown under normal to high light conditions. Cells were collected at mid-log phase from standard and high light conditions for total starch quantification by an Anthrone method. The color was measured with a spectrophotometer at 620nm. Three biological experiments were done (n=3), and error bars represent the mean ± SD (n = 4). The student *t*-test was calculated, and it significantly accepted (P < 0.0001) value indicated in the figure.

### Fatty acid composition under high light condition

LC/MS analysed the fatty acids transesterification and the resultant fatty acid methyl esters (FAMEs) from control strains *CC125, 137AH*, and mutants *pgrl1* and *pgr5* under normal (50 µmol photons m^−2^ s^−1^) and high light (500 µmol photons m^−2^ s^−1^) cells (Table 1). Different composition of fatty acids was observed in the high light grown *C. reinhardtii*. Predominant increase of Palmitic acid (C16:0), Palmitoleic acid (16:1), Oleic acid (C18:1) was observed along with a substantial amount of linoleic acid (C18:2) and linolenic acid (C18:3) under the high light condition in *pgrl1* and *pgr5* mutants. Significant increases in monounsaturated fatty acid were observed in *pgrl1* and *pgrl5* mutants under high light growth. The Palmitic acid (16:0) and Hexadecadienoic acid (16:2) increased to two folds under the high light condition in *pgr5*. However, these fatty acids were not changed much in the *wt, 137AH*, and *prgl1* cells. Further, the α-linolenic fatty acid (18:3) followed a similar trend increased as a two-fold level under high light conditions from *wt, 137AH, prgl1*, and *prg5*. In general, the α-linolenic acid is enriched with MGDG in the thylakoid membrane resulting in the α-linolenic acid exceeding under high light condition. Previous reports have proved that a high light intensity could enhance significantly Palmitic acid (C16:0), Stearic acid (C18:0), and Oleic acid (C18:1) content in *Pavlova lutheri*. Not many fatty acid profile changes were observed between low and high light of *CC125* and *137AH*. The monounsaturated fatty acid (MUFA) increased significantly in *pgrl1* and *pgr5* mutants with increasing light intensity. In contrast, a decrease in PUFA content was observed under the high light condition in *Pavlova lutheri* (Guedes *et al*., 2010). Further, the total fatty acid also increased in high light conditions. This result is in consistent with our TEM data that membrane damage occurred due to increased ROS under light conditions. This study shows that the monounsaturated fatty acid level was enhanced in *pgrl1* and *pgr5* mutant under 500 µmol photons m^−2^ s^−1^. In contrast, this fatty acid level was not changed much in normal light conditions.

**Table 1.** Fatty acid content of *C.reinhardtii* under normal and high light conditions. The data represents the mean n=3.

| Fatty acid | *CC125 50* | *CC125 500* | *137AH 50* | *137AH 500* | *pgrl1 50* | *pgrl1 500* | *pgr5 50* | *pgr5 500* |
| --- | --- | --- | --- | --- | --- | --- | --- | --- |
| **C14:0** | 0.9 53 ± 0.005 | 0.83 ±0.004 | 0.327 ± 0.002 | 0.666 ± 0.003 | 0.185 ± 0.001 | 0.176 ± 0.001 | 0.594 ± 0.003 | 0.609 ±0.003 |
| **C14:1** | 0.443 ± 0.002 | 0.3 ± 0.002 | 0.172 ± 0.001 | 0.599 ± 0.003 | 0.42 3 ± 0.002 | 0.154 ± 0.001 | 0.329 ± 0.002 | 0.508 ± 0.003 |
| **C14:2** | 0.62 ± 0.003 | 2.18 ± 0.011 | 0.386 ± 0.002 | 0.857 ± 0.004 | 0.423 ± 0.0002 | 0.132 ± 0.001 | 0.632 ± 0.003 | 0.406 ± 0.002 |
| **C14:3** | 1.507 ± 0.008 | 0.82 ± 0.004 | 0.105 ± 0.001 | 0.372 ± 0.0002 | 0.318 ± 0.002 | 0.64 ± 0.003 | 0.228 ± 0.001 | 0.216 ± 0.001 |
| **C16:0** | 2.618 ± 0.013 | 1.44 ± 0.007 | 1.392 ± 0.002 | 1.338 ± 0.002 | 2.123 ± 0.007 | 3.415 ± 0.007 | 3.45 ± 0.005 | 7.132 ± 0.006 |
| **C16:1** | 2.081 ± 0.022 | 3.21 ± 0.021 | 1.177 ± 0.006 | 2.491 ± 0.012 | 1.874 ± 0.004 | 4.178 ± 0.006 | 2.212 ± 0.003 | 4.398 ± 0.001 |
| **C16:2** | 0.088 ± 0.025 | 1.81 ± 0.009 | 0.262 ± 0.001 | 1.674 ± 0.003 | 0.36 ± 0.002 | 2.544 ± 0.003 | 0.573 ± 0.003 | 5.027 ± 0.005 |
| **C16:3** | 4.817 ± 0.0008 | 5.31 ± 0.027 | 1.102 ± 0.026 | 5.102 ± 0.009 | 1.816 ± 0.009 | 2.78 ± 0.014 | 2.1 ± 0.010 | 3.13 ± 0.016 |
| **C18:0** | 1.612 ± 0.013 | 2.19 ± 0.011 | 0.569 ± 0.003 | 1.037 ± 0.005 | 1.659 ± 0.008 | 2.061 ± 0.012 | 1.025 ± 0.005 | 3.752 ± 0.007 |
| **C18:1** | 2.557 ± 0.004 | 0.75 ± 0.004 | 0.171 ± 0.001 | 0.824 ± 0.004 | 0.439 ± 0.002 | 1.998 ± 0.004 | 0.826 ± 0.004 | 5.46 ± 0.012 |
| **C18:2** | 0.889 ± 0.034 | 0.68 ± 0.003 | 0.327 ± 0.002 | 0.213 ± 0.001 | 0.651 ± 0.003 | 0.212 ± 0.001 | 0.199 ± 0.003 | 1.15 ± 0.003 |
| **C18:3** | 1.808 ± 0.012 | 4.56 ± 0.109 | 3.015 ± 0.015 | 6.205 ± 0.036 | 2.725 ± 0.029 | 5.895 ± 0.014 | 5.738 ± 0.109 | 7.266 ± 0.046 |
| **C18:4** | 2.39 ± 0.013 | 3.97 ± 0.04 | 1.181 ± 0.006 | 3.376 ± 0.017 | 2.114 ± 0.011 | 1.23 ± 0.006 | 3.161 ± 0.04 | 3.395 ± 0.017 |
| **MUFA<sup>a</sup>** | 4.081 ± 0.037 | 5.26 ± 0.008 | 1.52 ± 0.008 | 3.361 ± 0.009 | 2.736 ± 0.027 | 5.33 ± 0.019 | 3.367 ± 0.011 | 10.366 ± 0.025 |
| **PUFA<sup>b</sup>** | 12.119 ± 0.11 | 24.6 ± 0.053 | 6.378 ± 0.058 | 18.79 ± 0.063 | 8.407 ± 0.203 | 12.43 ± 0.072 | 12.631 ± 0.042 | 20.59 ± 0.09 |
| **TFA<sup>c</sup>** | 22.383±0.173 | 28 ± 0.068 | 10.186 ±0.082 | 23.754 ± 0.085 | 15.11±0.252 | 25.415 ±0.101 | 21.067 ± 0.073 | 42.449± 0.131 |

<sup>a</sup> MUFA, monounsaturated fatty acids.
<sup>b</sup> PUFA, polyunsaturated fatty acids.
<sup>c</sup> TFA, total fatty acids.

## Discussion

Cyclic electron transport around PSI requires the functions of PGRL1 and PGR5 to generate a proton gradient over the thylakoid membrane. PGR5 plays a regulatory role in cyclic electron flow around PSI, which contributes indirectly to PSI protection by enhancing photosynthetic control, which is a pH-dependent down-regulation of electron transfer at the cytochrome *b*6*f* (Buchert *et al*., 2020). Less proton motive force across the thylakoid membrane and reduced cyclic electron transport around PSI suggest this protein’s pivotal role in the photosynthesis process to protect it from high light (Yadav *et al*., 2020). The cyclic electron transport is diminished in these mutants. It is known that cyclic electron transport plays a significant role in protecting the plants in light stress. Recently, we reported that the photosynthetic activity was severely affected in *pgrl1* and *pgr5* of *C. reinhardtii* when the cells were grown in high light (Yadav *et al*., 2020). We have also observed the non-photochemical quenching is substantially reduced in *pgr5*, which is supposed to protect the algae from high light that could harmlessly dissipate excess excitation energy as heat (Nama *et al*., 2019). The *pgr5* cells were much more sensitive to high light (500 µmol photons m^−2^s^−1^) when they grow in photoautotrophic conditions. To our knowledge, in *pgrl1* and *pgr5*, the autophagy induced lipid accumulation has not been explored from high light condition. This study shows that autophagy induces lipid accumulation under high light in *pgrl1* and *pgr5* mutant.

### High light effect the growth and chlorophyll content

Light is an essential factor in the production of biomass. Half of the microalgae species’ dry weight is carbon and lipid (Zhu *et al*., 2015). This study demonstrates that high light intensity (500 µmol photons m^−2^s^−1^) enhanced the growth rate, promoted biomass, and lipid accumulation of *C. reinhardtii* in comparison to lower light intensities (Figs. 1&5). *pgrl1* and *pgr5* genes proteins are essential in the acclimation process to high light; thus, here, the growth curves are evident that the absence of these leads to less biomass and growth. (Fig. 1A). The light stress response is previously studied, which shows the importance of light intensity in producing biomass and fatty acids in microalgae (Nzayisenga *et al*., 2020).

### High light induces the ROS and Autophagy in Chlamydomonas

On the other hand, when high light conditions are exposed to microalgal cells, PSII undergoes severe damage. Hence photosynthetic electron transport chain induces the ROS (Li *et al*., 2009). Antioxidant usually accumulates in cells like carotenoids (Perez-Martin *et al*., 2014)in order to repair the damage caused by ROS is observed in our results (Fig. S1). Interestingly, the ROS generation was much higher in *pgr5* because the lack of this gene led to reduced efficiency of photosynthesis. Also, since this mutant is also involved in the protection of PSI, which is also involved in the acceptor side limitation of PSI. Thus, there is a limitation of PSI’s acceptor side in *pgr5*, as one could expect more ROS generation at the PSI acceptor side (Arjun Tiwari *et al*., 2016). Further, lipid synthesis could serve as a receptor to excess electrons to acclimatize to the abiotic stress. Studies show that oxidative stress itself causes lipid accumulation under nitrogen stress in microalgae (Yilancioglu *et al*., 2014).

TEM images of Chlamydomonas showed a pronounced increase in vacuoles’ size under high light intensity, particularly in mutant strains (Fig. 6). It was reported in Chlamydomonas cells under nitrogen limiting conditions. They have proved that autophagy plays an essential role in lipid metabolism under nutrient limitation (Couso *et al*., 2016). It has been reported that in stress conditions (like oxidative stress, rapamycin stress, and ER stress), ATG8 gene was expressed in *C. reinhardtii* (Perez-Martin *et al*., 2014). In agreement with this hypothesis here we reported that significant expression of ATG8 protein was observed under high light condition by immunofluorescence microscopy, this indicates that ATG8 is a key protein to induce the autophagy process in Chlamydomonas (Fig. 3&4). An abundance of ATG8 protein was increased in mutant due to high light, especially in *pgr5*. It seems an increase in stromal redox poise, which induces autophagy, thereby lead to TAG accumulation. High light stress leads to ROS formation, which activates the autophagy mechanism. Increased ROS was seen in high conditions; however, this was more in *pgr*5 mutant when grown in high light that indicates that high light leads to an increase in oxidative stress and autophagy, which accompanying increased lipid content in the cell.

Autophagy of Chlamydomonas contains numerous autophagy-related proteins. Among various ATG proteins, ATG8 protein is essential for forming autophagosomes (Pérez-Pérez *et al*., 2016). In this study, we measured the expression level of ATG8 protein to evaluate the autophagy role in *137AH, pgrl1*, and *pgr5* (Fig. 3). ATG8 appeared as a faint band in the *wt* (*CC125* and *137AH*) and mutant (*pgrl1* and *pgr5*) grown under optimal condition (50 µmol photons m^−2^s^−1^), and it was increased when cells were shifted to high light. However, this increase is significant in *pgr5* mutant than *pgrl1*. Further, TEM confirmed several lytic vacuoles, and small vesicles inside the vacuoles were detected in high light treated cells. These changes could be due to ROS production (Fig. 2). As well as vacuoles size has been increased abundantly under high light in mutants, but the size of the cell has been reduced in *pgr5*. Our result supports that light stress leads to ROS. In turn, ROS induces autophagy, as reported earlier, leading to an increase in lipid production. Confocal and FACS data also suggested an increase in lipid droplet formation under high light in *pgr5* mutant (Fig. 5). Continuous light exposure leads accumulation of biomass and the cellular over reduction and formation of ROS, which induced autophagy and lipids in microalgae (Shi *et al*., 2017). Our previous results indicate that PSI and PSII were damaged when cells were grown in high light (Nama *et al*., 2019 and Yadav *et al*., 2020). Based on this result, we interpret that lipid accumulation in mutant strains could be explained that PSII or PSI is over reduce, which led to high ROS production. These increased ROS levels act as an autophagy inducer (Heredia-Martinez *et al*., 2018).

### High light induces the high amount of lipids

In microalgae, lipid and starch are the two primary energy storage forms and share the common carbon precursors for biosynthesis. Carbon partitioning is essential for the development of biofuels and chemicals. Most carbohydrates are stored as starch in microalgae. In several algal species, starch and lipid, both accumulated while in some species starch level decreases, and TAG increases, suggesting that lipid may be synthesized from starch degradation. Our results showed that both lipid and starch content increased in wild type and *pgrl1*. However, lipid accumulation precedes starch accumulation (Figs. 5 & 7), consistent with the previous studies in the microalga, *Pseudochlorococcum* under nitrogen stress condition (Li *et al*., 2011). However, the starch content was reduced in *pgr5* mutant under high light growth that explains that starch degradation would have been converted to lipids in *pgr5*. Complex metabolic regulations in the cell may control the accumulation of lipid droplets. The *pgr5* controls the proton gradient and PSI dependant cyclic electron transport; therefore, the altered NADPH and ATP ratio may switch the metabolic pathways during the photosynthesis process. Some of the metabolites, these metabolic regulation alterations, would have converted to lipids, which is necessary to acclimatize to the high light in *pgr5*. In *C. reinhardtii* mutant, which lacks cell walls, starch is the dominant sink for carbon storage. Rapid lipid synthesis occurs only when the carbon supply exceeds starch synthesis (Fan *et al*., 2012). In the model organism, *Arabidopsis thaliana* and Chlamydomonas TAG synthesis were originated from multiple types of acyltransferase. The final step of TAG biosynthesis was catalyzed by DGAT and PDAT, both involved in seed oil accumulation (Zhang *et al*., 2009). In the present study, we find the regulation of PDAT and DGTA at the protein level. Both the enzymes are accumulated under high light intensity, supporting this enzyme’s TAG synthesis (Fig. 3). Thus, the accumulation of TAG would be either DGAT or PDAT routes.

Interestingly, the thin layer chromatography results show that the membrane lipids have been decreased, which indicates that the degradation of these lipids could have converted to TAG, especially on *pgr5*. Similar reports were observed suggested that the recycling of membrane lipids, MGDG, and DGDG into TAG accumulation in *C. reinhardtii* (Abida *et al*., 2015). We assume that fatty acids are dissociated from glycolipids, which are involved in the Kennedy pathway to synthesize the TAGs through the DGAT enzyme (Yoon *et al*., 2012).

### High light alters the fatty acid composition in Chlamydomonas

The fatty acid composition also plays a significant role in biofuel production. Studies show that an increase in light intensity (300 µmol photon m^−2^ s^−1^) increases the percentage of C16:0 and C18:0 in *Neochloris oleoabundans* (Sun *et al*., 2014). *Pavlova luther*i also shows an increase in polyunsaturated fatty acids (PUFA) content under low light intensity (Freddy *et al*., 2014). The decrease in PUFA content shows the membrane system degradation, which shows that membrane lipids contain a high amount of UFA. In contrast, storage lipids contain more SFA in di flagellate due to photooxidation damage under high light, which resembles with the results in this study (Khanra *et al*., 2017). The *pgrl1* and *pgr5* mutants have a higher percentage of monounsaturated fatty acids (MUFA) than PUFA due to high light, as shown in Table 1. More number of PUFA is due to more alteration of lipid membranes when exposed to environmental conditions, which led to the formation of unsaturation in the lipids. C18:3 is one of the polyunsaturated fatty acids, significantly identified in the high light condition in *pgr*5 mutant than the other cell strains because of high light; it alters the thylakoid membranes disassociation of C18:3 fatty acid in the cell. Our results provided us complete evidence for the fact that the two mutant *pgrl1* and *pgr5* mutant of *C. reinhardtii* lead to the accumulation of fatty acid, biomass, and autophagy under high light conditions. The correlation between the stress and TAG accumulation in photosynthesis of algae has been known from a century ago. Under the high light condition, *pgrl1* and *pgr5* mutant fatty acid composition may be an essential study for biotechnological applications. Hence our study could offer an essential value that microalgae-based lipid production can be promoted by applying various strategies like feedstocks to biodiesel and animal feed.

## Acknowledgments

R.S was supported by the Department of Biotechnology (BT/PR14964/BPA/118/137/2015), *Council of Scientific and Industrial Research* (No.38 (1381)/14/EMR-II) and UGC-ISF Research Grant - File No. 6-8/2018 (IC), DST-FIST and UGC-SAP, Govt. of India, for financial support. We thank Gilles Peltier, CEA - CNRS - Aix Marseille Université, France, and Prof. Michael Hippler, University of Munster, Germany, for providing the mutants of *pgrl1*and *pgr5* and also critical reading of the manuscript.

## Supplementary Figure legends

**SFig. 1**. Total carotenoid content from the cells grown under normal and high light conditions. Carotenoids were calculated from cells grown with control *CC125*, and *137AH* and mutant *pgrl1* and *pgr5* cells grown from normal light (50 µmol photons m^−2^ S^−1^) and high light condition (500 µmol photons m^−2^ S^−1^) measured after 3^rd^ day. Three biological experiments were done, and values were presented with mean ±SD. (P < 0.0107).

**SFig. 2**. Separation of polar lipids by 2D-TLC from *Chlamydomonas reinhardtii* strains *CC125, 137AH, pgrl1*, and *pgr5* under normal and high light grown conditions. Dark spots indicate lipids. The expression profile was marked with an arrow.

**SFig. 3**. Neutral lipid content is quantified from the cells grown under normal to high light conditions. Neutral lipid content was qualitatively measured with Nile Red fluorescence from the cells *CC125, 137AH, pgrl1*, and *pgr5* grown with normal to high light condition stained with Nile Red on days 1–4. The measurements have been done with a microplate reader.

